# Isolation of ACE2-dependent and -independent sarbecoviruses from Chinese horseshoe bats

**DOI:** 10.1101/2023.03.02.530738

**Authors:** Hua Guo, Ang Li, Tian-Yi Dong, Hao-Rui Si, Ben Hu, Bei Li, Yan Zhu, Zheng-Li Shi, Michael Letko

**Author notes:** **Corresponding authors:** Michael Letko, Ph.D.,. Zheng-Li Shi, Ph.D. co-senior authors (Zheng-Li Shi, Michael Letko).

## Abstract

While the spike proteins from SARS-CoV and SARS-CoV-2 bind to host ACE2 to infect cells, the majority of bat sarbecoviruses cannot use ACE2 from any species. Despite their discovery almost 20 years ago, ACE2-independent sarbecoviruses have never been isolated from field samples, leading to the assumption these viruses pose little risk to humans. We have previously shown how spike proteins from a small group of ACE2-independent bat sarbecoviruses may possess the ability to infect human cells in the presence of exogenous trypsin. Here, we adapted our earlier findings into a virus isolation protocol, and recovered two new ACE2-dependent viruses, RsYN2012 and RsYN2016, as well as an ACE2-independent virus, RsHuB2019. Although our stocks of RsHuB2019 rapidly acquired a tissue-culture adaption that rendered the spike protein resistant to trypsin, trypsin was still required for viral entry, suggesting limitations on the exogenous entry factors that support bat sarbecoviruses. Electron microscopy revealed ACE2-independent sarbecoviruses have a prominent spike corona and share similar morphology to other coronaviruses. Our findings demonstrate a broader zoonotic threat posed by sarbecoviruses and shed light onto the intricacies of coronavirus isolation and propagation *in vitro*.

**SIGNIFICANCE:** Several coronaviruses have transmitted from animals to people and 20 years of virus discovery studies have uncovered thousands of new coronavirus sequences in nature. Most of the animal-derived sarbecoviruses have never been isolated in culture due to cell incompatibilities and a poor understanding of the *in vitro* requirements for their propagation. Here, we built on our growing body of work characterizing viral entry mechanisms of bat sarbecoviruses in human cells and have developed a virus isolation protocol that allows for exploration of these understudied viruses. Our protocol is robust and practical, leading to successful isolation of more sarbecoviruses than previous approaches and from field samples that had been collected over a 10-year longitudinal study.

## INTRODUCTION

With the increase of coronaviruses crossing the species barrier into humans and causing severe diseases over the last 20 years, significant effort has been invested into understanding coronaviruses in diverse animals, globally. The first viral relatives of SARS-CoV were discovered in Rhinolophus bats in 2005, demonstrating these animals are a natural reservoir for the sarbecovirus subgenus of the betacoronaviruses (1, 2). However, in comparison to SARS-CoV, these bat sarbecoviruses contained numerous polymorphisms in their spike glycoprotein – the viral protein responsible for binding cell receptor molecules and mediating viral invasion into host cells. Later, cell-culture based studies with these bat sarbecoviruses showed that although their spike proteins were not compatible with some human receptors, exchanging their spike genes with the SARS-CoV spike allowed for the viruses to replicate in cell culture - demonstrating that cell entry is a primary species barrier for bat sarbecoviruses (3). The identification of bat sarbecoviruses that could bind angiotensin-converting enzyme 2 (ACE2), the same receptor as SARS-CoV, has led to the overall assumption that bat sarbecoviruses which do not use this receptor pose little threat of zoonosis to humans.

In a broad screen of sarbecovirus entry, we found several host cell entry phenotypes that are determined by the presence or absence of deletions within the receptor binding domain (RBD) of the spike glycoprotein (4). Clade 1 RBDs do not contain any deletions and are capable of binding ACE2 from multiple species, clade 2 RBDs contain 2 deletions and do not use ACE2, clade 3 and 4 RBDs contain single deletion, but are capable of binding ACE2 more specifically from their host species (4-12). The first bat sarbecoviruses discovered were clade 2 viruses and any attempts to isolate them from field samples have failed (1, 2). We recently showed that a high concentration of trypsin could facilitate *in vitro* entry and replication of viral pseudotyped and recombinant sarbecoviruses containing clade 2 RBD spike proteins (4, 13). Many other viruses have been shown to replicate in the presence of trypsin, including several gastrointestinal coronaviruses with uncharacterized host receptors (14-17). Taken together, these findings suggest that some clade 2 bat sarbecoviruses may also have the capacity to infect human cells, which is a prerequisite for cross-species transmission to humans.

Here, we further optimized our methods for propagating clade 2 sarbecoviruses in culture for viral isolation from field samples. We successfully isolated one clade 2 RBD sarbecovirus as well as two new clade 1 RBD sarbecoviruses from *Rhinoliophus sinicus* fecal samples collected between 2012-2019, showing that the higher trypsin level used is compatible with both ACE2-dependent and -independent sarbecoviruses. Electron microscopy of virions showed that the spike density on clade 2 virions may vary from clade 1 RBD sarbecoviruses. This new sarbecovirus isolation protocol increases the chance of viral isolation from field samples and has extended our ability to explore and understand the biological features of less studied sarbecoviruses in the laboratory.

## RESULTS

### Isolation of three novel sarbecoviruses from Chinese horseshoe bats in the presence of trypsin

In our previous studies, we showed that some clade 2 sarbecoviruses are capable of entering and replicating in human cell lines in a high trypsin environment (4, 7, 13). To assess if trypsin-mediated entry is sufficient to support clade 2 virus isolation from field samples, we chose 18 bat rectal swabs or fecal samples from the WIV biobank, which were collected from individual bats during a seven-year longitudinal survey from 2012-2019. Sixteen of 18 samples tested positive for betacoronaviruses using an established reverse transcription (RT)-nested PCR targeting a fragment of the RNA-dependent RNA polymerase (RdRp) gene (**Supplementary table 1**) (18, 19). We also performed next-generation sequencing (NGS) on all 18 samples to obtain nearly full-length genome sequences for 15 viruses (**Supplementary table 1**), including two isolates, 7896 and 7909, which we have reported previously (5). In general, samples with lower Ct values produced sequence data, while samples with higher Ct values were somewhat less consistent in our NGS pipeline (**Supplementary table 1**). Based on our study of recombinant bat sarbecoviruses, we modified our virus isolation protocol to include a high concentration of trypsin, and a chilled centrifugation step during inoculation (see METHODS and (13)). With this modified protocol, we isolated three sarbecoviruses from positive samples, in a human liver cell line (Huh-7) and have named them: RsYN2012 (sample 4105), RsYN2016 (sample 162173) and RsHuB2019 (sample 190366) (**Fig. 1A**). We further examined the genome sequence of the three isolates and found that they shared a similar genome structure and organization with other bat and human sarbecoviruses (**Fig. 1B**). Based on the RBD portion of the spike that we and others have previously used to group sarbecoviruses into clades, RsYN2012 (4105) and RsYN2016 (162173) belong to clade 1, and RsHuB2019 (190366) belongs to clade 2 (**Fig. 1A-C**). Comparing whole genomes, the two clade 1 viruses were 99.9% and 98.3% similar to bat SARS-related CoV, RsWIV1, while the clade 2 virus RsHuB2019 (190366) showed 93.2% nucleotide similarity with bat SARS-related CoV, HKU3-1 (**Fig. 1B, Table 1**). All three viruses were only approximately 80% similar to SARS-CoV-2, and less than 80% of similar with clade 3 and 4 viruses. (**Fig. 1B, Table 1**). The largest sequence variation for any of our isolates was in the spike gene of the clade 2 virus, RsHuB2019 (190366), which exhibited only between 65-77% similarity to the clade 1 virus spikes genes (**Fig. 1B, Table 1**).

**Table 1.** Genomic comparison of viral isolates with other sarbecoviruses. Genomic comparison of RsHuB2019, RsYN2012, and RsYN2016 with SARS-CoV, SARS-CoV-2 and their related CoVs

| Sequence identities with SARS-CoV, SARS-CoV-2 and related bat CoVs (nt/aa %) |  |  |  |  |  |  |  |  |  |  |  |  |
| --- | --- | --- | --- | --- | --- | --- | --- | --- | --- | --- | --- | --- |
|  |  | Full-length genome | ORF1a | ORF1b | S | ORF3 | E | M | ORF6 | ORF7a | ORF7b | N |
| RsHuB2019<br>(190366) | SARS-CoV | 88.3 | 88.2/94.4 | 92.5/99.0 | 76.4/79.5 | 83.0/82.9 | 97.4/100 | 94.6/97.3 | 93.2/90.6 | 93.8/91.9 | 93.3/93.3 | 97.2/97.9 |
|  | SARS-CoV-2 | 79.6 | 76.0/80.9 | 86.1/95.5 | 73.1/77.9 | 75.3/74.2 | 95.2/96.1 | 84.2/90.1 | 75.3/66.1 | 84.2/86.9 | 84.1/77.3 | 88.7/91.0 |
|  | Bat SARSr-CoV RsWIV1 | 88.6 | 88.3/94.2 | 92.5/99.1 | 76.4/79.3 | 83.5/83.6 | 97.8/100 | 95.2/98.2 | 92.2/89.1 | 93.5/95.1 | 93.3/95.6 | 96.6/97.6 |
|  | Bat SARSr-CoV HKU3-1 | 93.2 | 94.0/97.3 | 93.2/98.7 | 85.5/89.2 | 95.3/95.3 | 100/100 | 98.5/98.6 | 98.4/96.9 | 94.9/96.7 | 97.0/100 | 96.9/97.2 |
|  | Bat SARSr-CoV BM48-31 | 78.6 | 76.8/81.6 | 85.6/96.1 | 70.2/74.4 | 71.6/68.6 | 90.5/92.2 | 80.5/91.0 | 66.1/52.4 | 63.3/58.0 | 60.2/63.4 | 78.6/88.0 |
|  | Bat SARSr-CoV RaTG15 | 74.5 | 71.3/76.5 | 83.6/94.5 | 65.9/68.9 | 70.1/66.9 | 85.3/81.8 | 79.5/92.3 | 67.8/55.9 | 63.4/54.9 | 51.1/33.3 | 78.7/88.5 |
| RsYN2012<br>(4105) | SARS-CoV | 95.6 | 96.9/97.9 | 96.4/99.3 | 90.2/92.4 | 98.5/97.1 | 99.1/100 | 97.3/98.2 | 95.2/92.2 | 93.0/92.7 | 93.3/93.3 | 98.5/99.8 |
|  | SARS-CoV-2 | 79.6 | 76.0/80.5 | 85.9/95.6 | 73.9/78 | 75.9/74.2 | 95.6/96.1 | 84.8/90.1 | 78.0/72.6 | 85.5/88.5 | 84.1/77.3 | 88.5/91.0 |
|  | Bat SARSr-CoV RsWIV1 | 99.9 | 100/100 | 100/100 | 100/99.9 | 99.8/99.6 | 100/100 | 100/100 | 100/100 | 92.7/95.1 | 93.3/95.6 | 100/100 |
|  | Bat SARSr-CoV HKU3-1 | 88.2 | 88.1/94.2 | 90.9/98.6 | 77.7/80.1 | 83.0/82.5 | 97.8/100 | 94.0/96.8 | 93.2/89.1 | 96.2/96.7 | 97/100 | 96.1/96.4 |
|  | Bat SARSr-CoV BM48-31 | 78.9 | 77.0/81.4 | 85.7/96.2 | 70.9/75.8 | 73.4/72.7 | 90.9/92.2 | 80.3/90.5 | 65.1/52.4 | 65.3/58.8 | 60.2/63.4 | 78.5/88.0 |
|  | Bat SARSr-CoV RaTG15 | 74.5 | 71.2/76.5 | 83.5/94.5 | 65.6/70.1 | 70.5/66.5 | 85.7/100 | 78.9/91.4 | 66.7/62.7 | 64.8/56.6 | 51.1/33.3 | 79.0/88.0 |
| RsYN2016<br>(162173) | SARS-CoV | 95.6 | 97.0/98.2 | 96.5/99.4 | 90.2/92.5 | 97.6/96.4 | 99.6/100 | 97.0/98.2 | 94.8/92.2 | 94.6/94.3 | 95.6/93.3 | 98.6/99.8 |
|  | SARS-CoV-2 | 79.7 | 76.0/80.9 | 86.3/95.7 | 73.7/77.9 | 75.6/73.8 | 94.3/96.1 | 84.8/90.1 | 78.0/72.6 | 84.7/87.7 | 85.6/81.8 | 89.0/91.4 |
|  | Bat SARSr-CoV RsWIV1 | 98.3 | 98.8/100 | 98.3/99.8 | 96.5/98.8 | 98.3/98.2 | 98.7/100 | 99.7/100 | 99.5/100 | 99.7/99.2 | 100/100 | 98.7/99.5 |
|  | Bat SARSr-CoV HKU3-1 | 88.2 | 88.3/94.5 | 90.9/98.7 | 77.5/80.2 | 82.5/81.8 | 97.0/100 | 93.7/96.8 | 92.7/89.1 | 93.0/95.9 | 92.6/95.6 | 96.5/96.9 |
|  | Bat SARSr-CoV BM48-31 | 78.8 | 77.2/81.6 | 85.6/96.3 | 71.1/75.8 | 73.3/72.0 | 90.0/92.2 | 80.5/90.5 | 65.1/52.4 | 64.1/58.8 | 63.4/68.3 | 78.9/88.5 |
|  | Bat SARSr-CoV RaTG15 | 74.6 | 71.2/76.8 | 83.6/94.6 | 65.4/69.9 | 70.2/65.8 | 85.3/81.8 | 78.9/91.4 | 66.1/62.7 | 63.4/54.9 | 52.6/35.6 | 79.1/88.5 |

**Figure 1.**
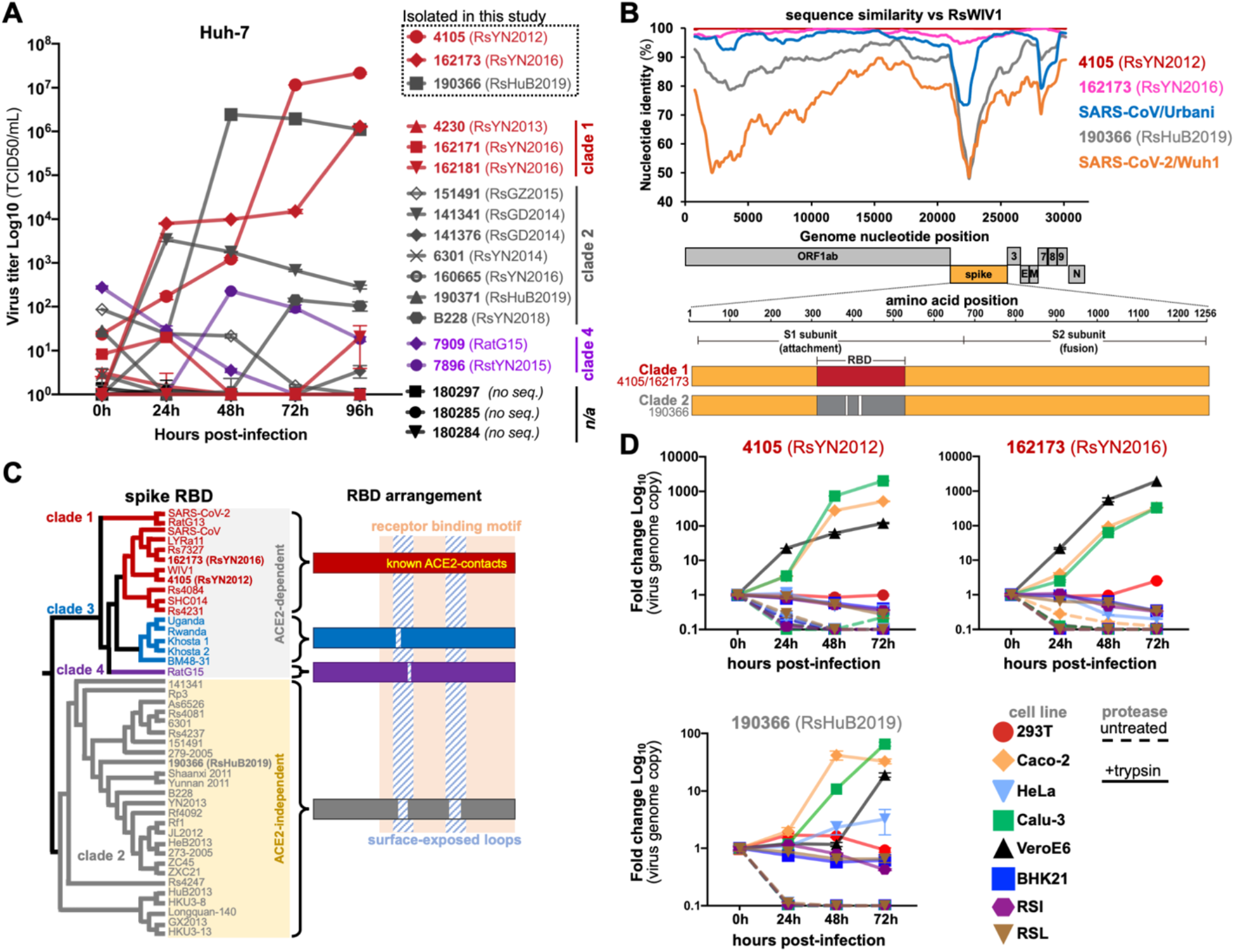
Isolation of clade 1 and clade 2 RBD sarbecoviruses on human cell lines. (**A**) Field samples were used to inoculate Huh-7 cells in the presence of trypsin. Viral titers were quantified in supernatants by qRT-PCR. (**B**) Whole genome nucleotide sequences were compared to RsWIV1 with a sequence similarity plot. Open reading frame (ORF) positions indicated under the X-axis.(**C**) Cladogram analysis of RBD amino acid sequences (corresponding to SARS-COV spike 323-510) for sarbecoviruses. RBD indels and receptor preferences are indicated for each functional phylogenetic clade. Viruses isolated in this study are in bold font. (**D**) Viral isolates were inoculated on indicated cell cultures and viral replication was monitored by qRT-PCR.

### Cellular tropism of the three bat sarbecovirus

To assess if the three bat sarbecoviruses pose similar cellular or tissue tropism with known sarbecovirus and further assess their risk of interspecies transmission, we conducted infectivity assays in several cell lines common in coronavirus research. We found, in addition to Huh-7 cells, the two clade 1 viruses, RsYN2012 (4105) and RsYN2016 (162173), could replicate efficiently in human cell lines (Caco-2, Calu-3), and African Green Monkey cells (VeroE6) in the presence of trypsin, but poorly infected these cells in the absence of trypsin (**Fig. 1D**). The clade 2 virus, RsHuB2019 (190366), could replicate in Caco-2, Calu-3, and VeroE6 cells, similar to the clade 1 viruses, although with lower entry and replicate efficiency (**Fig. 1D**). In addition, HeLa cells were semi-permissive to clade 2 virus RsHuB2019 (190366) infection in the presence of trypsin (**Fig. 1D**). Consistent with our prior study, both clade 1 and 2 virus were unable to replicate efficiently in baby hamster kidney (BHK-21), and two bat primary cell lines, including *Rhinolophus sinicus* (*R*.*s*.) intestine (RSI) and lung (RSL), in the presence or absence of trypsin (**Fig. 1D**)(13).

### ACE2 is the receptor for RBD clade 1 but not RBD clade 2 sarbecovirus

To explore the receptor usage of the three-novel bat sarbecoviruses, we performed virus infectivity studies using BHK-21 cells expressing known coronavirus receptors from humans and bats. Consistent with prior studies (4, 6, 13), we found that only the clade 1 virus could utilize human ACE2 for cell entry and that the clade 2 virus, RsHuB2019 (190366) could not use any known coronavirus receptor, with or without trypsin (**Fig. 2A**).

**Figure 2.**
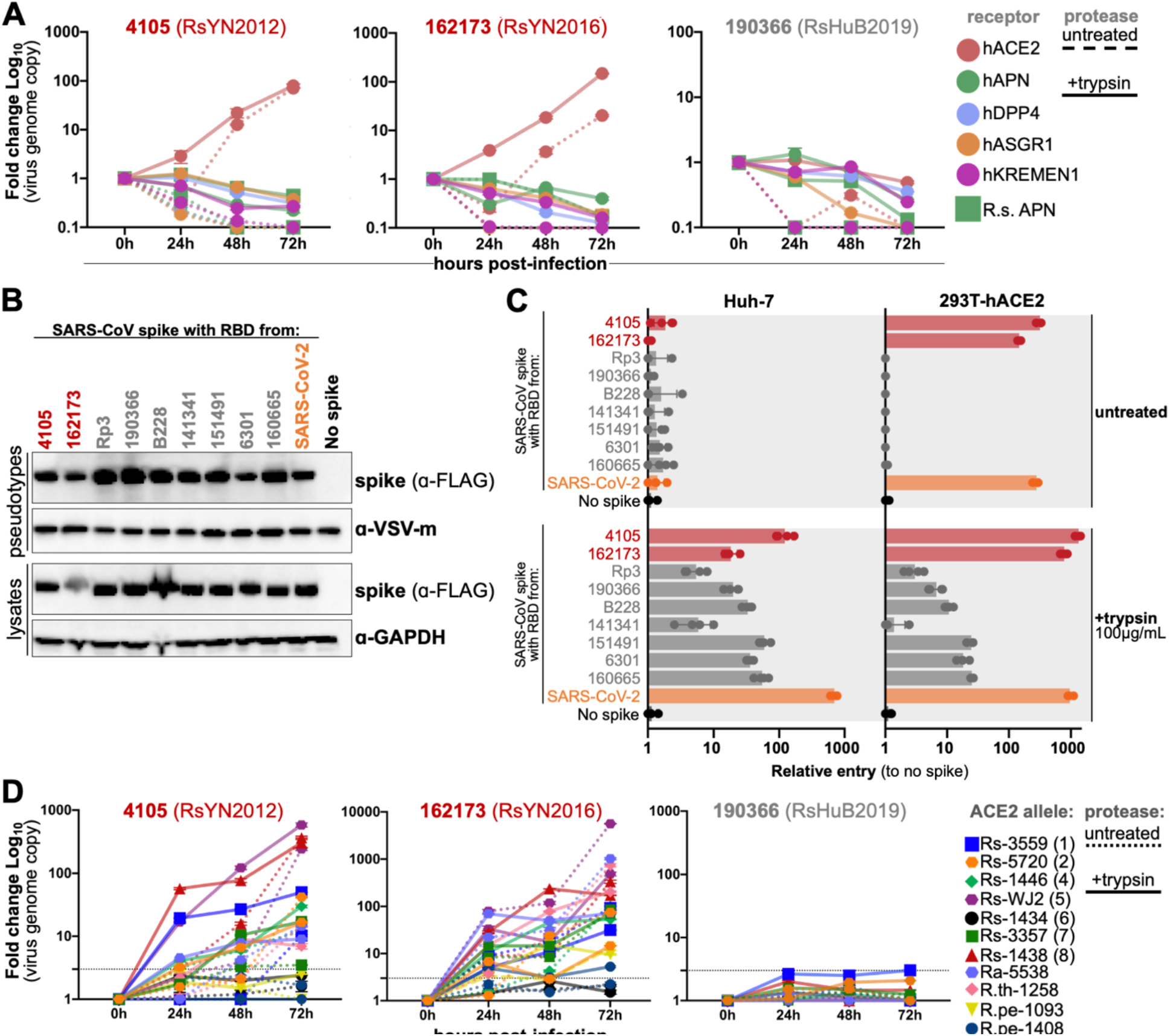
Clade 2 RBD sarbecoviruses do not use any known coronavirus receptors for cell entry. (**A**) BHK cells were transfected with human orthologues of known coronavirus receptors and then infected with viral isolates. Replication was quantified by qRT-PCR. (**B**) VSV pseudotyped bearing chimeric SARS-CoV spikes with the indicated virus RBDs were generated in HEK 293T cells and concentrated in OptiPrep. Spike was detected in cell lysates and pseudotyped by probing for FLAG. (**C**) Huh-7 cells or cells transduced to express human ACE2 were infected with viral pseudotyped and luciferase was measured as a readout for cell entry. (**D**) BHK cells were transfected with the indicated bat ACE2 alleles and infected with viral isolates. Replication was monitored by qRT-PCR.

To assess the cell entry capacity of the viruses we failed to isolate from the other samples, we assembled a panel of recombinant RBD chimeras, with SARS-CoV chimeric spike containing the RBD sequence from the different samples (4). Of the eight clade 2 virus-positive samples we attempted to isolate virus from, four samples contained newly identified viruses (B228, 190366, 141341, 151491), while the RBD sequences in the other samples were identical to other RBDs from this study or RBD sequences we have previously tested (Rs4081, As6526; **Supplementary table 1**) (4, 5, 7, 13). The RBDs for clade 1 viruses 4105 and 162173 are identical to RBDs from RsWIV1 and Rs7327, respectively, which we have also previously tested (4). For comparison, we included a SARS-CoV spike chimera with the RBD from SARS-CoV-2 and a clade 2 RBD from the prototypical virus, Rp3 (**Fig. 2B-C**) (10). All RBD chimeras exhibited similar levels of incorporation into VSV pseudotyped particles (**Fig. 2B**). We have previously shown exogenous trypsin allows mediated sarbecovirus entry into otherwise poorly susceptible cell lines, Huh-7 and 293T (4, 7, 13). Transduction of 293T cells with human ACE2 allows for clear detection of ACE2-dependent entry (10, 11). Consistent with the live virus infection assay results, only pseudotyped with clade 1 virus RBDs were capable of entering and transducing human ACE2 expressing cells without trypsin, but not any of the clade 2 viruses (**Fig. 2C**). As we have shown for other clade 2 RBDs, the addition of trypsin dramatically increased entry for these spikes, with a notable exception for the RBD from sample 141341 (**Fig. 2C**).

Previous studies from our groups and others have reported that the ACE2 gene is diverse across bat species (12, 20-23). We have shown that the ACE2 gene is highly polymorphic in Chinese horseshoe bats (*R. sinicus)*, and that different ACE2 alleles within the same species exhibit different susceptibility to various sarbecoviruses infection (20). To further confirm if the ACE2 orthologues from different bat species or different Chinese horseshoe bat (*R. sinicus)* ACE2 alleles support the entry of clade 2 viruses, we tested a large panel of bat ACE2 alleles for their ability to support live virus infection in BHK-21 cells. Consistent with our previous study (20), we found the clade 1 virus, RsYN2012 (4105) and RsYN2016 (162173), could utilize most alleles from *R. sinicus* ACE2, as well as ACE2 from *Rhinolophus affinis* and *Rhinolophus thomasi*, for cell entry regardless of trypsin (**Fig. 2D**). RsYN2016 (162173) could also enter the BHK-21 cell expressing *Rhinolophus pearsonii* (R.pe) ACE2-1093 with low efficiency, but not the allele 1408. However, RsYN2012 (4105) could not use either of the ACE2 alleles from *Rhinolophus pearsoni* for entry (**Fig. 2D**). In contrast, we found none of these bat ACE2 genes supported replication of clade 2 virus RsHuB2019 (190366), even in the presence of trypsin (**Fig. 2D**). Taken together, these results demonstrate that only clade 1 viruses we isolated possess the capacity to use the ACE2 from different species and that the clade 2 virus employs an unknown molecule(s) for entry in human cells that is distinct from other coronaviruses.

### Tissue culture adaptations in ACE2-independent spike increase trypsin resistance

Coronavirus spike genes are known to rapidly acquire cell-culture specific adaptations – sometimes in as few as three passages (24-31). Over the course of this study, we replenished our viral stocks by subsequently passaging the previous stock in Huh-7 cells, leading to three viral passages (experiments from figure 1 are passage 1, figure 2 are passage 2 and figure 4 are passage 3). We extracted viral RNA from the remainder of each stock after each passage and looked for potential cell-culture adaptations, across the whole viral genome by next-generation sequencing. We found one nonsynonymous (T24550G) substitution that emerged at low frequency in the clade 2 virus, RsHuB2019 (190366) at the 1^st^ passage, resulting in V976L mutation in the S gene. By the third passage, we observed an increase in the frequency of spike V976L mutation (from 61% to 99.4%) with L976 becoming the dominant polymorphism (**Fig. 3A**). We did not observe additional mutations elsewhere in the RsHuB2019 genome or in the genomes of the clade 1 RBD viruses, RsYN2012 (4105) and RsYN2016 (162173).

**Figure 3.**
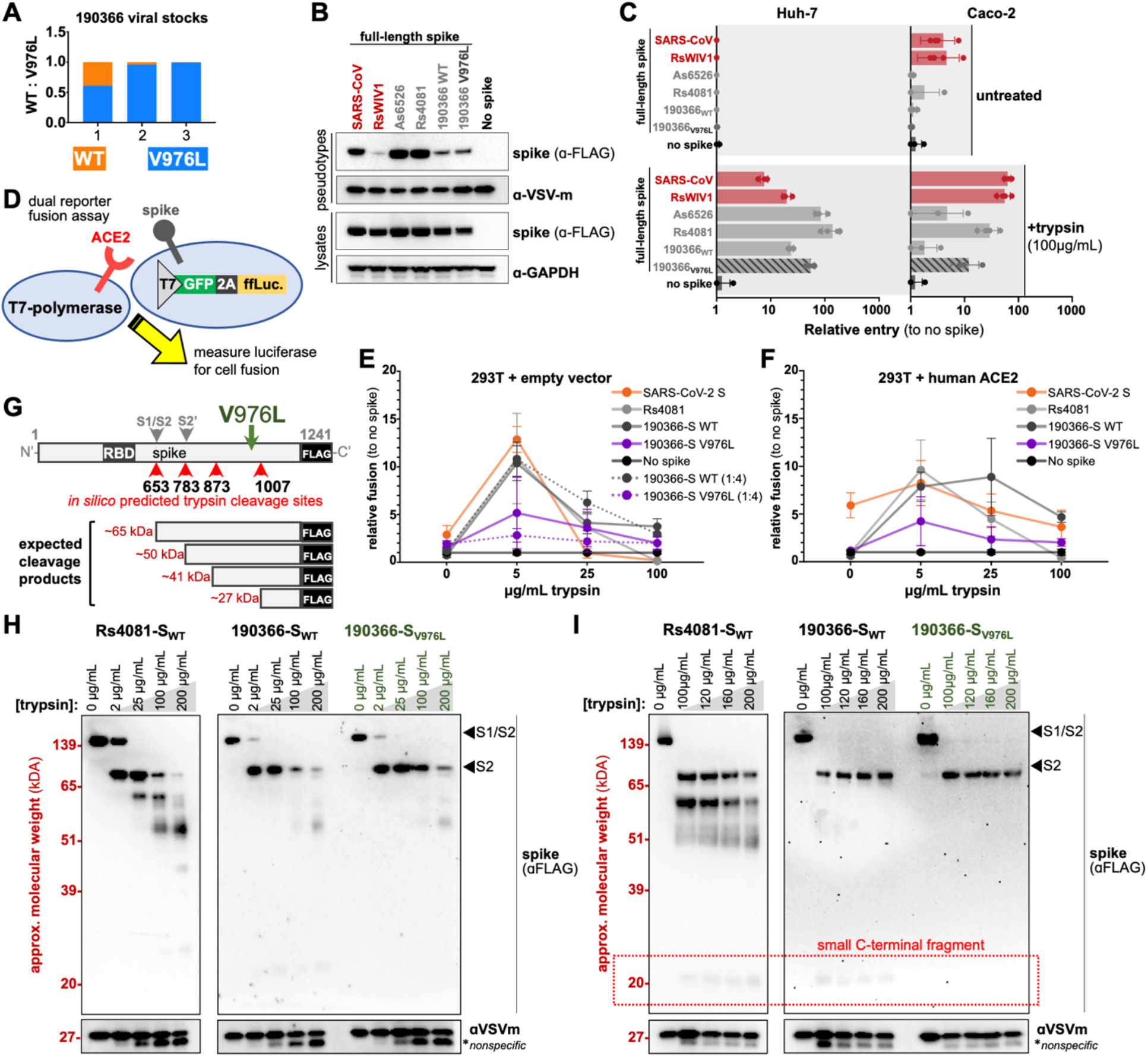
Clade 2 RBD virus adaptation to cell-culture. (**A**) V976L mutation emerged in 190366 virus stocks. (**B**) Pseudotyped particles were produced with full-length spike WT or the V967L mutant. Spike was detected in producer cells and pseudotyped by western blotting for FLAG. (**C**) Indicated cells were infected with viral pseudotyped in the presence or absence of trypsin. (**D**) Schematic overview of the dual-reporter fusion assay developed for this study. T7 polymerase drives expression of GFP and luciferase separated by a P2A fusion peptide. (**E**) HEK 293T cells expressing receptor or, (**F)** empty vector and T7-polymerase were combined with cells expressing spike and the T7-driven reporter. Luciferase was measured as a readout for cell fusion. Dotted lines indicate data from 1:4 ratio of receptor:spike cells. (**G**) Overview of 190366 spike with *in silico* predicted trypsin digest sites indicated. Location of V976L is indicated in green. (**H**) Concentrated viral pseudotyped were combined with a wide range or trypsin dilutions or (**I**) a fine range of trypsin dilutions and incubated at 37°C. Spike digestion was assessed by western blot for FLAG epitope.

**Figure 4.**
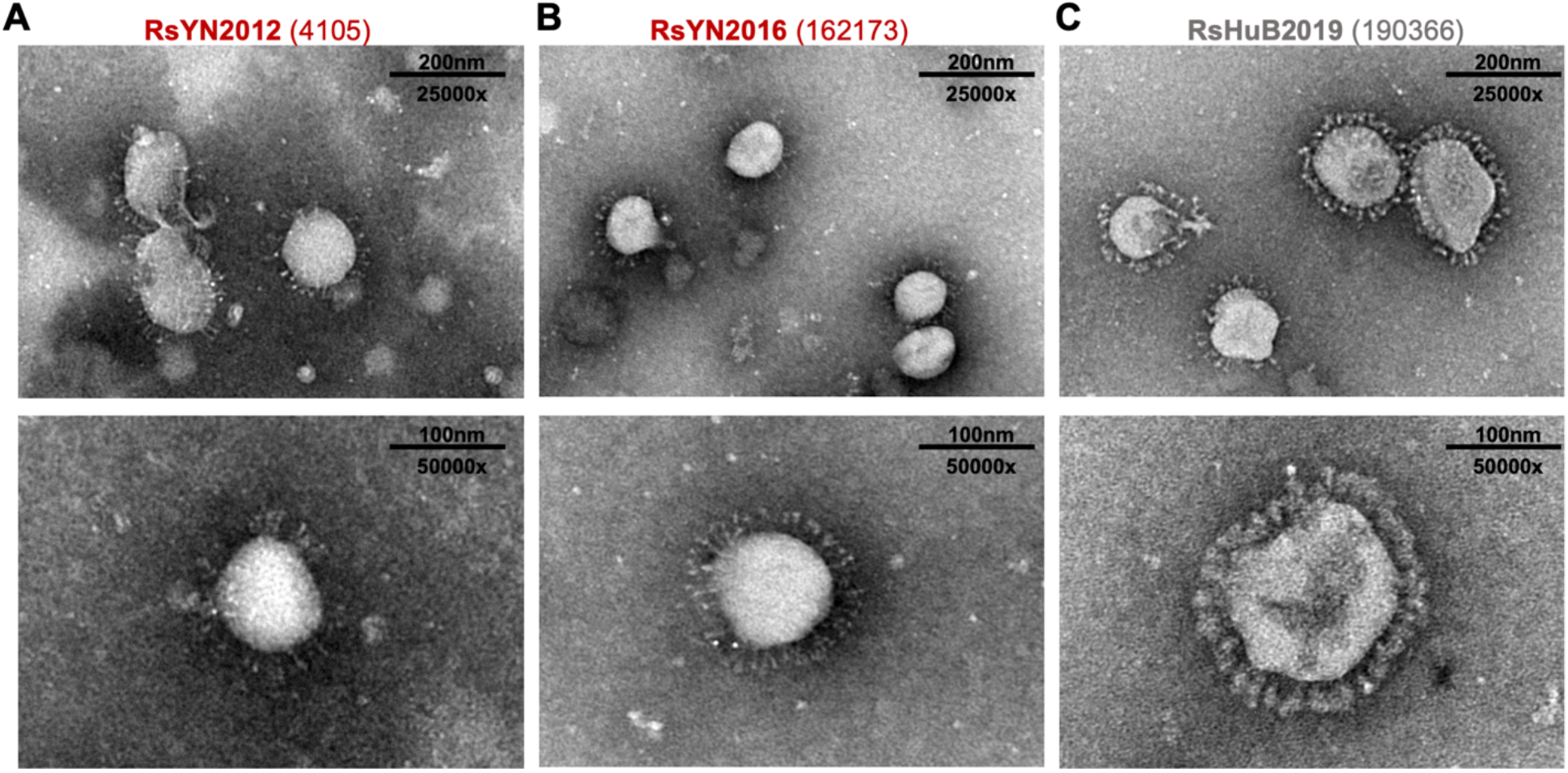
Electron microscopy of purified viral isolates. Viral stocks for. (**A**) RsYN2012 (**B**) RsYN2016 or (**C**) RsHuB2019 were visualized by transmission electron microscopy. Bottom images were taken at a higher magnification to show detail.

To characterize the V976L mutation in the RsHuB2019 (190366) spike, we constructed VSV-based pseudotyped containing full-length spike with either V976 or L976 and tested their cell entry in Huh-7 and Caco-2 cell lines. Full-length spike from clade 1 viruses, SARS-CoV, and RsWIV1, as well as clade 2 viruses, Rs4081 and As6526, were used as comparative controls (13) (**Fig. 3B-C**). We found that V976L mutation did not increase spike incorporation into virions (**Fig. 3B**), but moderately enhanced the entry of RsHuB2019 (190366) in both Huh-7 and Caco-2 cell lines, only in the presence of trypsin (**Fig. 3C**).

Because the V976L mutation is in close proximity to the host cell fusion machinery present in the spike S2 domain, we wondered if this mutation was influencing the fusogenic properties of RsHuB2019 spike. To test if this mutation modulated spike cell fusion properties, we performed a cell-cell fusion assay similar to previous approaches by combing cells individually expressing spike or receptor and a complementary reporter system (32). HEK 293T cells expressing T7 polymerase and human ACE2 or empty vector were combined, 1:1, with HEK 293T cells expressing a T7-driven reporter cassette and spike (**Fig. 3D**). Because RsHuB2019 spike had reduced incorporation into pseudotyped (**Fig. 3B**), we also included a condition with 4 times the amount of spike containing cells to receptor cells (**Fig. 3E**; *1:4, dotted line*). Increasing the concentration of trypsin to even 5 μg/mL resulted in more than a 10-fold increase in cell fusion for spikes with clade 1 and clade 2 RBDs, while the addition of ACE2 to the cells increased basal entry of SARS-CoV-2 spike without trypsin (**Fig. 3E, F**). Curiously, regardless of the ratio between spike-expressing cells and target cells, viral fusion was reduced for the RsHuB2019 spike with V967L mutation compared to the wildtype spike (**Fig. 3E, F**; *dotted lines*).

To further explore how the V976L mutation in increased spike cell entry in the presence of trypsin, we tested the *in vitro* trypsin resistance of spike, with the clade 2 virus Rs4081 as control.Purified V976 or L976 pseudotyped particles were combined with different amounts of trypsin, incubated at 37°C for 5 min, and spike degradation was analyzed by western blot. As we have previously shown, trypsin cleaved the Rs4081 spike into several fragments, including the expected fragments corresponding to cleavage at the S1/S2 boundary as well as a secondary, S2’ site, at 25μg/mL or above trypsin (13) (**Fig. 3G, H**). In contrast, RsHuB2019 spike displayed less of these degradation products, with the V976L mutation showing resistance to trypsin digestion at 100μg/mL - the concentration we used to propagate virus in our cultures (**Fig. 3H**). When we performed a second trypsin digestion between 100-200 μg/mL and used a more sensitive western blot substrate, a smaller digestion product, the approximate size of a C-terminal fragment of spike that is predicted to digest from a site near V976, was absent from the V976L mutant but present for Rs4081 and WT RsHuB2019 spike (**Fig. 3G, I; boxed in red**). Thus, V967L may reduce trypsin digestion in spike near this mutation. Taken together, these findings strongly suggest the clade 2 virus spike adapted to the exogenous (porcine) trypsin included during viral propagation, rather than the cell lines themselves.

### Electron microscopy of purified virions reveals potential difference between RBD clades

In order to confirm if we had isolated the three sarbecorviruses successfully, we purified viral stocks over a 30% sucrose cushion and processed the samples for analysis by transmission electron microscopy. Purified viral particles displayed typical coronavirus morphology under electron microscopy: virions were approximately 100–120 nm in diameter, with “corona-like” ring of spike glycoproteins at the surface. Interestingly, the glycoprotein layer on clade 2 virions appeared denser than on clade 1 RBD virions (**Fig. 4A-C, S1A-C**).

## DISCUSSION

Although hundreds of sarbecoviruses have been discovered in animals, more than two-thirds of these viruses have clade 2 RBD spikes, which contain indel mutations that prevent them from using host ACE2 as a cell receptor (2-4, 6, 11, 12). Attempts to isolate these ACE2-independent sarbecoviruses from field samples have failed, hampering downstream laboratory-based assessments and leading to the general assumption that they pose little threat to humans. However, we have demonstrated the RBDs from a small group of these viruses are capable of mediating human cell entry, which we have verified with whole spike proteins and most recently, complete sarbecovirus replication recovered through reverse genetics (4, 7, 13). Here, we developed a virus isolation protocol built on these findings that is suitable for recovering both ACE2-dependent and –independent sarbecoviruses from bat fecal samples, underscoring the broader zoonotic threat posed by sarbecoviruses and the complexities underlying coronavirus cell entry.

Our successful isolation of a clade 2 RBD sarbecovirus (RsHu2019) and two clade 1 RBD sarbecoviruses: RsYN2012 (4105) and RsYN2016 (162173) suggest our approach is broadly applicable for sarbecoviruses, and an improvement over existing sarbecovirus isolation protocols. Notably, we isolated a viable virus (RsYN2012) from a field sample that had been in storage for more than 10 years (**Supplementary table 1**). The viruses we isolated were from samples with some of the lowest Ct values of the samples tested, suggesting higher viral titers are ideally required for successful isolation (**Supplementary table 1**). The only clade 2 RBD sample with a lower Ct value (141341) than the sample we isolated from (190366) also contained a viral RBD that was the least compatible with human cell entry, providing one explanation for why we failed to recover virus from this sample (**Supplementary table 1, Fig. 2C**).

The ACE2-dependent viruses we isolated, RsYN2012 and RsYN2016, were strikingly similar two other sarbecoviruses we have previously isolated or tested: RsWIV1 and Rs7327 (**Fig. 1B, Table 1** (4, 19). RsWIV1 and RsYN2012 were collected from the same location and time during the same sampling mission, which likely explains this close similarity (**Supplementary table 1**). However, the high similarity observed between RsWIV1 and viruses collected at later time points, including RsYN2016, suggests evolutionary constraints on these viruses in their hosts.

Coronaviruses acquire mutations when grown in cell culture and can rapidly adapt to the conditions and cells used for their propagation (24-31). Sequencing the viral stocks produced for this study revealed the emergence of a tissue-culture adaptation in the clade 2 virus, RsHuB2019, which appeared to increase viral entry in pseudotype experiments (**Fig. 3A-C**). Because this mutation was in close proximity to known fusion machinery in the S2 region of spike, we assessed spike fusion in a standard molecular assay and observed this mutation actually reduced fusion efficiency compared to wild-type spike (**Fig. 3D-F**). A close inspection of western blots following trypsin treatment of concentrated pseudotyped particles revealed this mutation resulted in the loss of a low-molecular weight digestion product, suggesting the mutation enhanced spike resistance to the trypsin used in our protocol (**Fig. 3G-I**). The trypsin we used in our studies is porcine-derived and not TPCK-treated, which may allow for additional spike digestion compared to TPCK-treated trypsin. Importantly, RsHuB2019 spike V967L still required trypsin for entry into cells (**Fig. 3C**), suggesting that the clade 2 viruses may not be capable of readily “evolving away from” trypsin dependence. Thus, while our protocol is suitable for the isolation of sarbecoviruses, more studies are needed into the species-specific proteases utilized by these viruses, which may lead to further protocol changes that reduce the development of cell-culture adaptations.

Electron microscopy of clade 1 and clade 2 virus isolates revealed a potential difference in the spike corona surrounding each virion. The spike trimers on ACE2-dependent clade 1 viruses appeared thinner and less evenly distributed than clade 2 virions, which may help explain clade 1’s increased sensitivity to trypsin versus clade 2 viruses (**Fig. 4, S1**). The virus stocks used for electron microscopy contained trypsin at the time of processing, therefore the differences in the fullness of the spike corona may reflect the overall trypsin resistance we have previously noted for the clade 2 RBD spikes (13). As the virus stocks used in our electron microscopy are from a later passage, we cannot exclude the possibility that this distinction may also derive from the presence of spike mutation V976L in RsHuB2019.

Other betacoronaviruses may provide clues about the entry mechanisms for clade 2 sarbecoviruses. For example, the bat merbecoviruses, PDF2180 and neoCoV, contain RBD deletions that prevent them from using host dipeptidyl peptidase IV (DPP4) as their receptor, and have been shown to require trypsin for their cell entry and propagation in human cell cultures (15). However, a recent study has shown these viruses bind to host ACE2 as a receptor and that providing this receptor can effectively remove the protease requirement (33). While we and others have shown the clade 2 sarbecoviruses do not use any known coronavirus receptor, our studies strongly suggest these viruses do rely on a conserved host molecule for entry (**Fig. 2**) (4, 6, 7, 11-13). Thus, more studies are needed to identify the receptor for clade 2 sarbecoviruses. Taken together our viral isolates demonstrate a cluster of bat sarbecoviruses that can infect human cells using mechanisms distinct from known human sarbecoviruses.

## METHODS

### Cells

HEK 293T, HEK 293T/17, BHK-21, VeroE6, Calu-3, and Hela were obtained from the American Type Culture Collection (ATCC), Caco-2 was generously gifted by Prof. Qin-Xue Hu. Bat-derived cell lines RSI and RSL were stored at the Wuhan Institute of Virology according to the previously described (13, 34). HEK 293T, HEK 293T/17, BHK-21, VeroE6, Huh-7, and Hela were maintained in Dulbecco’s modified Eagle medium (DMEM) supplemented with 10% fetal bovine serum (FBS). Calu-3, RSI, and RSL were maintained in Dulbecco’s Modified Eagle Medium/Nutrient Mixture F-12 supplemented with 15% fetal bovine serum (FBS). Cultures were maintained at 37°C with 5% CO2. All cell lines used in this study were species verified by cytochrome sequencing and tested negative for mycoplasma contamination by PCR as described previously (4, 35).

### Plasmids

Expression plasmids for human ACE2, human DPP4, human APN, human ASGR1, human KERMEN1, *Rhinolophus sinicus* APN, *Rhinolophus affinis* ACE2, and different alleles of *Rhinolophus sinicus* ACE2, were described previously (13, 20). *Rhinolophus pearsonii* ACE2-1093 and 1408, *Rhinolophus thomasi* ACE2, were amplified from the bat intestine as described previously (20).

The spike or RBD coding sequences for SARS-CoV/-2, RsWIV1, Rs4081, As6526, Rp3, 4105, 163173, B288, 141341, 151491, 190366, and 190366-S-V976L were codon optimized for human cells as previously described (13). Plasmid encoding T7-promoter driven dual reporter GFP and luciferase was generated by cloning firefly luciferase downstream of GFP in pUC19-T7-IRES-GFP. pUC19 - T7 pro - IRES - EGFP was a gift from Fei Chen (Addgene plasmid # 138586; http://n2t.net/addgene:138586; RRID: Addgene_138586). All the plasmids used in this study were verified by Sanger sequencing.

### Virus isolation

Bat fecal swabs or fecal samples were collected from several provinces in China over a seven-year period and stored at −80°C as described previously (19). The bat species was confirmed by cytochrome b sequence analysis as described previously (19). For virus isolation, the fecal samples were thawed on ice and centrifuged at 10,000g for 10 min at 4 °C before use. The supernatant (in 200 μL buffer) was filtered through 0.45 μm membranes and diluted 1: 2 with cold DMEM, trypsin was added to a final concentration of 625 μg/mL. Trypsin used for virus propagation was standard cell culture grade 0.25% porcine trypsin with EDTA and phenol red (ThermoFisher). Huh-7 cells were seeded on a 24-well plate and incubated at 37 °C overnight, then washed by DMEM once before incubating with 300 μL trypsin-treated samples. Inoculated plates were centrifuged at 1,200 g at 4 °C for 1 h, then incubated at 37 °C overnight. Approximately 20-24 hours post-infection, the monolayer cells were supplied with 300 μL fresh DMEM plus 4% FBS to a final concentration of 2% FBS and continued to incubate at 37 °C for 96 h. Cell-free supernatant was collected daily and detected for the presence of virus by RT–PCR.

### Pseudotyped virus production and entry assay

The coronavirus spike pseudotyped entry assays were performed as previously described with minor adjustments (4, 7, 10, 13). In brief, target cells were seeded in a 96-well plate and washed with PBS once before inoculating with equivalent volumes of pseudotyped stocks in the presence or absence of trypsin. Inoculated plates were centrifuged as described above. Entry efficiency was quantified 18-20 hours post-transduction, by measuring the luciferase activity using Bright-Glo luciferase reagent (Promega), following manufacturers’ instructions. Relative entry was calculated as the fold-entry in relative luciferase unit over the no spike control. All experiments were performed at least three times in triplicate.

### Cell-cell fusion assay

HEK 293T cells were seeded in a 6-well format. One group of cells was transfected with equivalent amounts of human ACE2 plasmid or empty plasmid and T7-polymerase plasmid. The second group of cells was transfected with equivalent amounts of spike expression plasmid and the dual reporter construct. 24 hours post-transfection, cells were trypsinized, diluted to 1×10^6^ cells/mL, and combined in either 1:1 or 1:4 ratios (receptor: spike transfected cells). 24 hours post-combining, cells were washed in cold PBS, and the cell culture media was replaced with trypsin-media and subsequently centrifuged at 1200 g at 4 degrees for 1 hour (to mimic the spin-infection procedures used in the infection assays). 24 hours post trypsin treatment and centrifugation, luciferase was measured on a plate reader using the bright-glo luciferase reagent (Promega).

### Electron Microscope Imaging

Virion concentration, purification and negative staining were performed as previously described with minor adjustments (19). In brief, fresh virus stocks were harvested at 72 hours post-infection, then centrifuged at 5,000 g for 30 min at 4 °C. Cell-free supernatants were collected and fixed by 0.1% formaldehyde at 4 °C overnight. Inactivated virions in the supernatant were loaded onto 5 ml of 30% sucrose in PBS buffer and centrifuged at 25,000 rpm in the SW28 rotor at 4°C for 2.5 hours. The pelleted virions were suspended in cold PBS, then applied to the grids and stained with 2% phosphotungstic acid (pH 7.0) on ice. The specimens were examined using a Tecnai transmission electron microscope (FEI) at 200 kV. Images were taken at a magnification of 25,000× and 50,000×.

### Phylogenetic analysis

Routine sequence management and analysis were carried out using DNAStar. Sequence alignments were created by Clustal W method in MegAlign from DNAStar package with default parameters. Maximum Likelihood trees with sarbecovirus spike RBD amino acid sequences were generated using PhyML 3.0 (36) with 1000 bootstrap replicates (37) and visualized as a cladogram in FigTree v1.4.4 (https://github.com/rambaut/figtree), as previously described (4, 10). Sequence similarity plot was generated using whole genomes for RsWIV1, SARS-CoV/Urbani, SARS-CoV-2 and isolates from this study using Simplot with the Kimura model, a window size of 1500 base pairs and a step size of 150 base pairs. (GenBank accession number: KF367457.1, AY278741.1, NC_045512.2).

### Viral replication detected by real-time RT–PCR

To study viral replication, target cells were seeded in a 24-well-plate and washed with DMEM once before inoculating with virus stocks in the presence or absence of trypsin. For receptor usage assays, BHK-21 cells were transfected with plasmids expressing different receptors 18-20 hours before infecting by the authentic virus with or without trypsin treatment. The inoculated plates were centrifuged at 1200 g at 4 °C for 1 hours and continued to incubate in a 37 °C incubator for 72 h. Cell-free supernatants (50 μL each time) were collected at 0, 24, 48 and 72 hours post-infection and stored at -80 °C for future use. Viral RNA was extracted and subjected to RT-PCR as previously described (13). Viral replication was quantified by RT-PCR using primers targeting the RdRp gene, forward primer: 5′-TTGTTCTTGCTCGCAAACATA-3′;reverseprimer:5′-CACACATGACCATCTCACTTAA-3′. The RNA from RsWIV1 stocks with known titers was used as a standard control to correlate the CT value and virus titer of the other viruses. All samples were analyzed in duplicate on two independent runs. One representative dataset is shown.

### Western blot

To check for cell expression of spike, HEK 293T cells producing viral pseudotyped were lysed in 1% SDS lysis buffer, clarified by centrifugation and blotted for FLAG as described previously (4). To check for spike incorporation, viral-like particle stocks were concentrated over a 10% Opti-Prep cushion in PBS at 21000 g for at 4°C 2 h, and blotted for FLAG on a 10% Bis-Tris gel (ThermoFisher) (4). Spike degradation was measured as in Guo et al. 2022, whereby concentrated pseudotyped stocks were incubated with trypsin concentrations at 37°C for 5 min, boiled, and blotted for FLAG (13). The substrate used in figure 3H is SuperSignal Western Blot Substrate Pico (ThermoFisher), and for increased sensitivity in figure 3I: SuperSignal Western Blot Substrate Atto (ThermoFisher).

### Statistical analysis and graphing

All graphed data are three technical replicates that are representative of at least 3 biological replicates. Graphed data was analyzed and visualized in GraphPad Prism version 9.

### Data availability

The nearly full-length genome sequences of SARSr-CoVs obtained in this study have been deposited in the GenBank database and the accession numbers are OQ503495-506, respectively. The accession number of *Rhinolophus pearsonii* ACE2-1093 and 1408, *Rhinolophus thomasi* are OQ511289-291.

### Biosafety and biosecurity

Laboratory work with VSV pseudotyped in mammalian cell lines was performed according to standard operating procedures (SOPs) under biosafety level 2 (BSL2) conditions that were approved by institutional biosafety committees (IBC) at WSU and Wuhan Institute of Virology (WIV). Work with bat SARS-related CoV was approved by the WIV IBC for this SOP and conform to the recommended guidelines for animal coronaviruses listed in the 6^th^ edition of Biosafety in Microbiological and Biomedical Laboratories (BMBL)(38). WIV facilities for this work adhere to the safety requirements recommended by the China National Accreditation Service for Conformity Assessment.

## Acknowledgments

We thank the core facility of the Wuhan Institute of Virology for their technical support. We also thank Pei Zhang and Ding Gao from the core facility of the Wuhan Institute of Virology for their help with the ultracentrifugation and Electron Microscopic Analysis. Work performed at WIV was jointly supported by the strategic priority research program (XDB29010101 to Z-LS), Key project (2020YJFK-Z-0149 and KJZD-SW-L11 to Z-LS) of the Chinese Academy of Sciences, National Natural Science Foundation of China (31727901 and 31770175 to Z-LS), National Key R&D program of China (2022YFC2305101 to BH) and work from the Paul G. Allen School for Global Health was supported by Washington State University.

## Author Contributions

M.L., H.G. and Z.-L.S. conceived and designed the study. H.G. performed virus isolation. H.G. and A.L performed virus infection experiments and electron microscopic analysis. M.L. performed pseudotyped experiments. M.L. developed the fusion assay and performed fusion experiments. T.-Y.D., H.-R.S., B.L., and Y.Z. performed the NGS and analyzed the data. B.H. performed Simplot analysis. H.G. and M.L. collected and analyzed data, and assembled figures. M.L., H.G. and Z.-L.S. wrote the manuscript.

## SUPPLEMENTARY INFORMATION

**Supplementary table 1.**
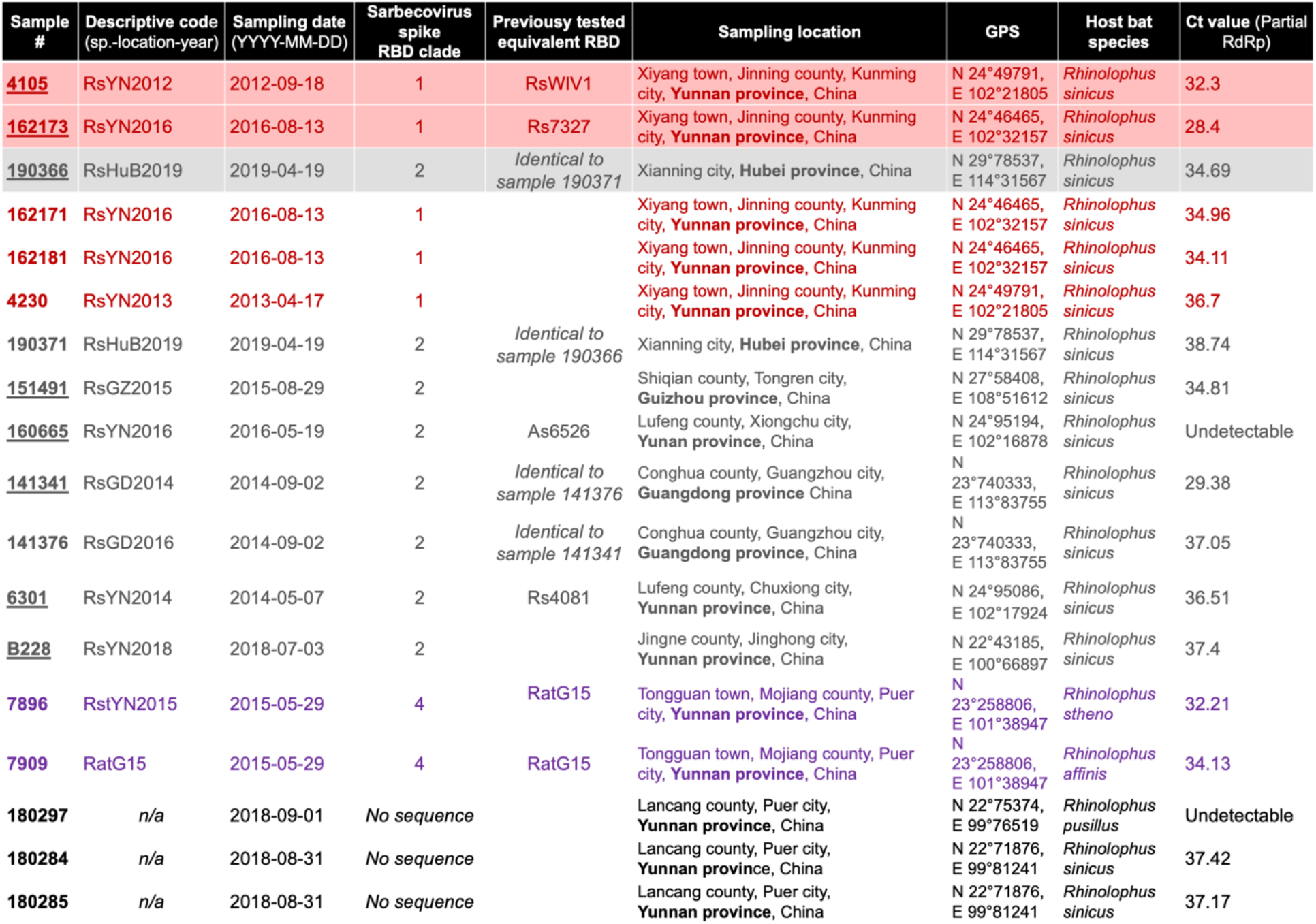
Metagenomic information regarding the samples from this study with isolated viruses in highlighted rows and viruses for pseudotyped experiments underlined

**Supplementary figure 1.**
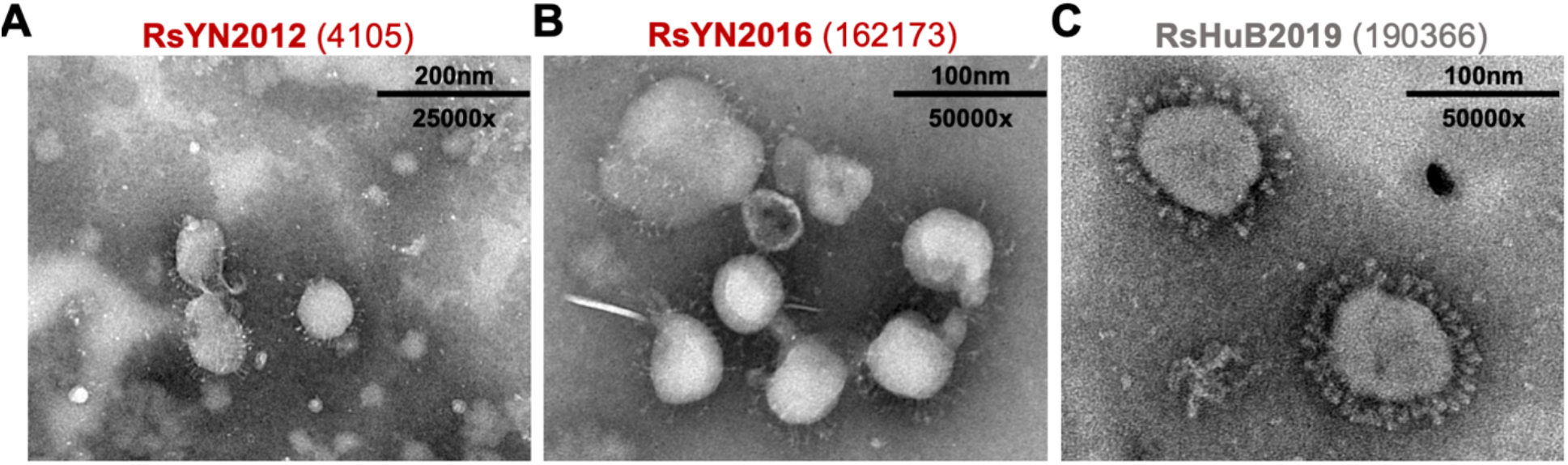
Additional electron micrographs of purified viral isolates. (**A**) RsYN2012 (**B**) RsYN2016 or (**C**) RsHuB2019 were visualized by transmission electron microscopy.

